# Studentsourcing - aggregating and re-using data from a practical cell biology course

**DOI:** 10.1101/2023.10.09.561479

**Authors:** Joachim Goedhart

## Abstract

Practical courses mimic experimental research and may generate valuable data. Yet, data that is generated by students during a course is often lost as there is no centrally organized collection and storage of the data. The loss of data prevents its re-use. To provide access to these data, I present an approach that I call studentsourcing. It collects, aggregates and reuses data that is generated by students in a practical course on cell biology. The course runs annually, and I have recorded the data that was generated by >100 students over 3 years. Two use cases illustrate how the data can be aggregated and re-used either for the scientific record or for teaching. As the data is obtained by different students, in different groups, over different years, it is an excellent opportunity to discuss experimental design and modern data visualization methods such as the superplot. The first use case demonstrates how the data can be presented as an online, interactive dashboard, providing real-time data of the measurements. The second use case shows how central data storage provides a unique opportunity to get precise quantitative data due to the large sample size. Both use cases illustrate how data can be effectively aggregated and re-used.

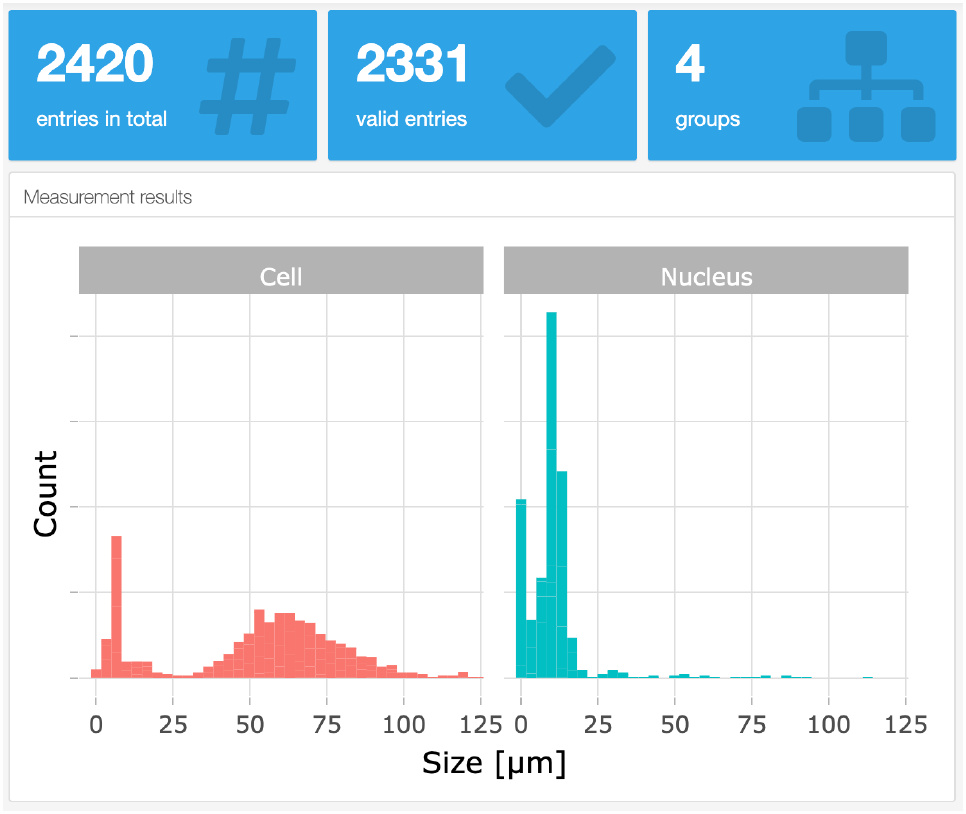

## Introduction

Teaching practical skills in a lab course is a crucial part of education in biology, biomedical science, and life sciences (Hofstein and Lunetta 2003; Reid and Shah 2007). In these lab courses data is generated, reported, and interpreted, much like real experimental lab work. However, students use their data just for their own lab report and the data is not centrally stored or aggregated. Consequently, most of the data that is gathered in a lab course is lost. Yet, these data are potentially useful. Especially for larger course, an impressive amount of data under well-controlled conditions can be generated. Therefore, by collecting and aggregating the data of multiple students over multiple years, one can easily gather a large dataset with high numbers of independent observations (Lazic, Clarke-Williams, and Munafò 2018).

Microscopy is an essential tool in cell biology. The use of microscopes to observe cells and organisms has changed from a qualitative, descriptive approach, into a quantitative method (Senft et al. 2023; Renz 2013; Wait, Reiche, and Chew 2020; Waters 2009). The development of digital cameras and image analysis software has catalyzed this transition (Carpenter 2007). Therefore, experiments that use microscopes are often followed by bioimage analysis to extract quantitative information from the data. To teach these skills, we combine a basic course on microscopy with teaching image processing and analysis in ImageJ/FIJI (Schneider, Rasband, and Eliceiri 2012). In a typical year, over one hundred students are enrolled in this course and therefore, a substantial amount of data is generated in the course.

I decided to collect the data that was generated by the students and store the measurement results in a central location. The data by itself can be valuable for the scientific community as precise estimates with good statistics can be obtained. Moreover, the data are a starting point to discuss data visualization, experimental design and how experimental design affects the statistics and interpretation of data. Here, I report the methods to collect, process and visualize the data. The data re-use is demonstrated in two use cases.

## Methods

For full reproducibility, this document is written using Quarto (Posit, https://quarto.org/), and the source code of the manuscript and the notebooks, and the data are availble in a repository: https://github.com/JoachimGoedhart/MS-StudentSourcing. A version rendered as HTML is avalaible and it provides easy access to the notebooks as well: https://joachimgoedhart.github.io/MS-StudentSourcing/

For full reproducibility, this document is written using Quarto (Posit, https://quarto.org/), and the source code of the manuscript and the notebooks, and the data are available in a repository: https://github.com/JoachimGoedhart/MS-StudentSourcing. A version rendered as HTML is available and it provides easy access to the notebooks as well: https://joachimgoedhart.github.io/MS-StudentSourcing/

The use cases presented here are part of a practical course that runs annually at the University of Amsterdam as part of the BsC programme “Biomedical Sciences”. In a typical year ∼120 students are enrolled and they are randomly assigned to four different groups (A/B/C/D) that take the course at different days. The students perform the experiments in pairs. Except in 2021 when, due to COVID-19 regulations, the students did the experiments individually.

### Use case 1

#### Sample preparation and measurements

A buccal swab is used to harvest cheek cells by scraping ∼5 times over the inside of the cheek. The tip of the sample collector is dipped into an eppendorf tube with 40 µl PBS, and the cells are transferred to an object slide by touching the slide with the tip. Next, 10 µl of 0.1% methyleneblue solution is added and the sample is enclosed by a square #1 coverslip (22 × 22 mm). The sample is used immediately for observation.

#### Microscopy

A Leica DM750 microscope with a manual XY-stage, equipped with a Lumenera Infinity 2-1RC CCD camera (1392 × 1040, 4.65 µm square pixels) was used for observation. Samples were illuminated with a LED and observed in transmitted light mode. A Leica Hi Plan 20× (NA 0.40) or 40× (NA 0.65) objective is used to observe the cells. Images were acquired using the Infinity Capture software. A separate image of a micrometer (Electron Microscopy Sciences 6804208, Stage Micrometer S8, Horizontal Scale, 1 mm Length) is acquired at the same magnification. The images are processed in ImageJ/FIJI (Schneider, Rasband, and Eliceiri 2012) and the dimensions of the images are calibrated with the micrometer image (Figure 1). The line tool is used to measure the diameter of the cells (the longest axis).

**Figure 1:**
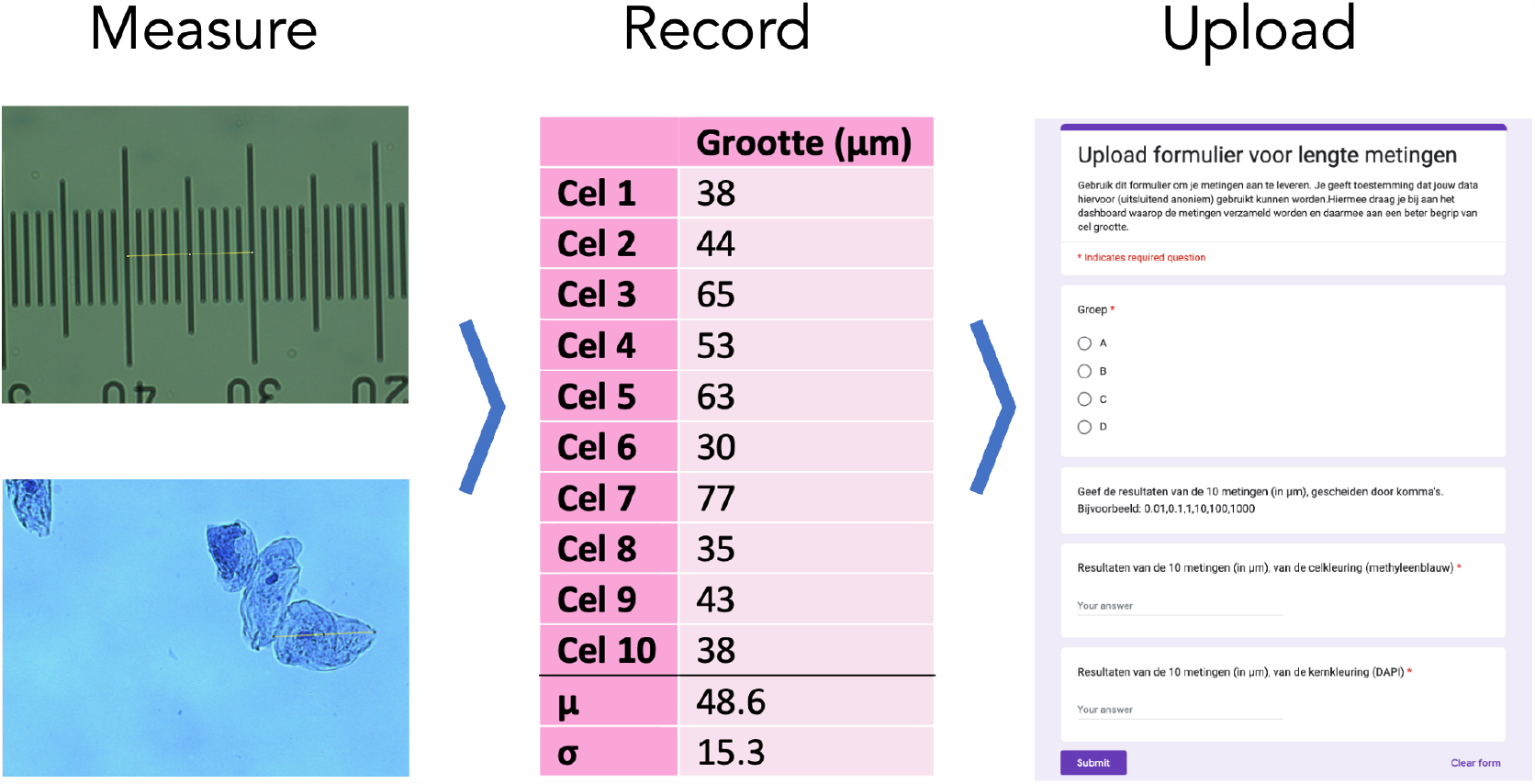
Overview of steps taken by the students to acquire and upload their data. The size of cells is measured by microscopy, the data is processed and recorded in their lab journal and finally, the data are uploaded by a Google form

#### Data collection

The data of the measurements is collected through a Google Form, an example of which is shown in Figure 1. By submitting the form, the students give permission for the anonymous use of the data. The data that is recorded by the form is the group (A/B/C/D), the size measurements of the cheek cells and the size measurements of the nucleus. The data is aggregated in a Google Sheet which has four columns with data on Timestamp, Group, size of cells, size of nuclei. When correctly uploaded, the two columns with the size data have comma separated values of 10 measurements.

#### Data processing

The data that is in the Google Sheet can be downloaded and read into R as a CSV. All subsequent processing, data visualisation, and presentation in dashboard style is done in R. The code is available on Github: https://github.com/JoachimGoedhart/CellSizeR. The cleaning of the data consists of removing empty cells, changing the column names, listing all individual measurements in a single row, forcing the data into a ‘numeric’ type and filtering for sensible values (anything outside the generous range of 0-1000 will be removed). A detailed protocol that explains the processing is available as protocol 10 (Goedhart 2022).

#### Data visualisation

A dashboard is composed in R Markdown with the {flexdashboard} package. The code is available here: https://github.com/JoachimGoedhart/CellSizeR and the live dashboard is available online: https://amsterdamstudygroup.shinyapps.io/CellSizeR/

### Use case 2

#### Sample preparation and measurements

HeLa cells are cultured according to standard procedures and seeded 1 or 2 days before the treatment on 12 mm diameter #1 glass coverslips. HeLa cells are incubated with 10 µM EdU (5-Ethynyl-2’-deoxyuridine, Lumiprobe) for 30 minutes at 37 °C. The cells are fixed with 4% formaldehyde in PBS and permeabilised with 0.1% Triton X-100 in PBS. Click chemistry is performed with 9 µM Cy3-azide (Lumiprobe) and 2 mM CuSO_4_. To start the reaction, 20 mg/ml ascorbate (final concentration) is added and the solution is used immediately to stain the cells. After 30 minutes, the cells are washed 3x with PBS and the sample is incubated with 0.1 µg/ml DAPI for 5 minutes. Samples are mounted in Mowiol (24% glycerol (w/v), 0.1 g mL–1 polyvinylalchol 4.88, 0.1 M Tris-HCl (pH8.5) in H2O) and used for observation with fluorescence microscopy.

#### Microscopy

A Leica DM750 microscope with a manual XY-stage, equipped with a Lumenera Infinity 2-1RC CCD camera (1392 × 1040, 4.65 µm square pixels) and a 100W HBO mercury lamp was used for observation. A Leica Hi Plan 20× (NA 0.40) or 40× (NA 0.65) objective is used to observe and image the cells. Images of at least 100 cells are acquired with DAPI (excitation 350/50nm, dichroic mirror 400nm, emission >420nm) and TRITC (excitation 540/25nm, dichroic mirror 570nm, emission >590nm) filters sets using the Infinity Capture software. The nuclei in both channels are counted by hand, or using an automated method, i.e. by segmentation and ‘particle analysis’ in imageJ to calculate the percentage of cells that are positive for Cy3 fluorescence, reflecting cells in the S-phase.

#### Data collection

The data of the measurements is collected through a Google Form. By submitting the form, the students give permission for the anonymous use of the data. The data that is recorded is the group (A/B/C/D), the percentage of cells in the S-phase for two methods, i.e. manual and using ImageJ/FIJI. The form is easy to set up and the data is collected in Google Sheets, yielding four columns; Timestamp, Group, and two columns with percentages of S-phase determined by the two methods.

#### Data processing & visualization

The data that is in the Google Sheet can be downloaded and read into R (R Core Team 2022) as a CSV. All subsequent processing and data visualization is done with R and quarto. The cleaning of the data consists of removing empty cells, changing the column names, conversion to a tidy format, forcing the data into a ‘numeric’ type and filtering for sensible values (anything outside the generous range of 0-100 will be removed).

## Results

### Use case 1: Comparing new results with historical data

The aim of the experiment is to determine the average size (diameter) of a human cheek cell and nucleus. To this end, the students acquire images of their own, stained cheek cells and measure the size of the cell and its nucleus. At least 10 measurements are made and the data are uploaded with a Google form. The average from each sample is an independent observation as it originates from a unique human specimen. To evaluate the accuracy of their own measurements, the students can compare their data with the historical data that is displayed on an online, interactive dashboard: https://amsterdamstudygroup.shinyapps.io/CellSizeR/. A snapshot of the dashboard is shown in Figure 2.

**Figure 2:**
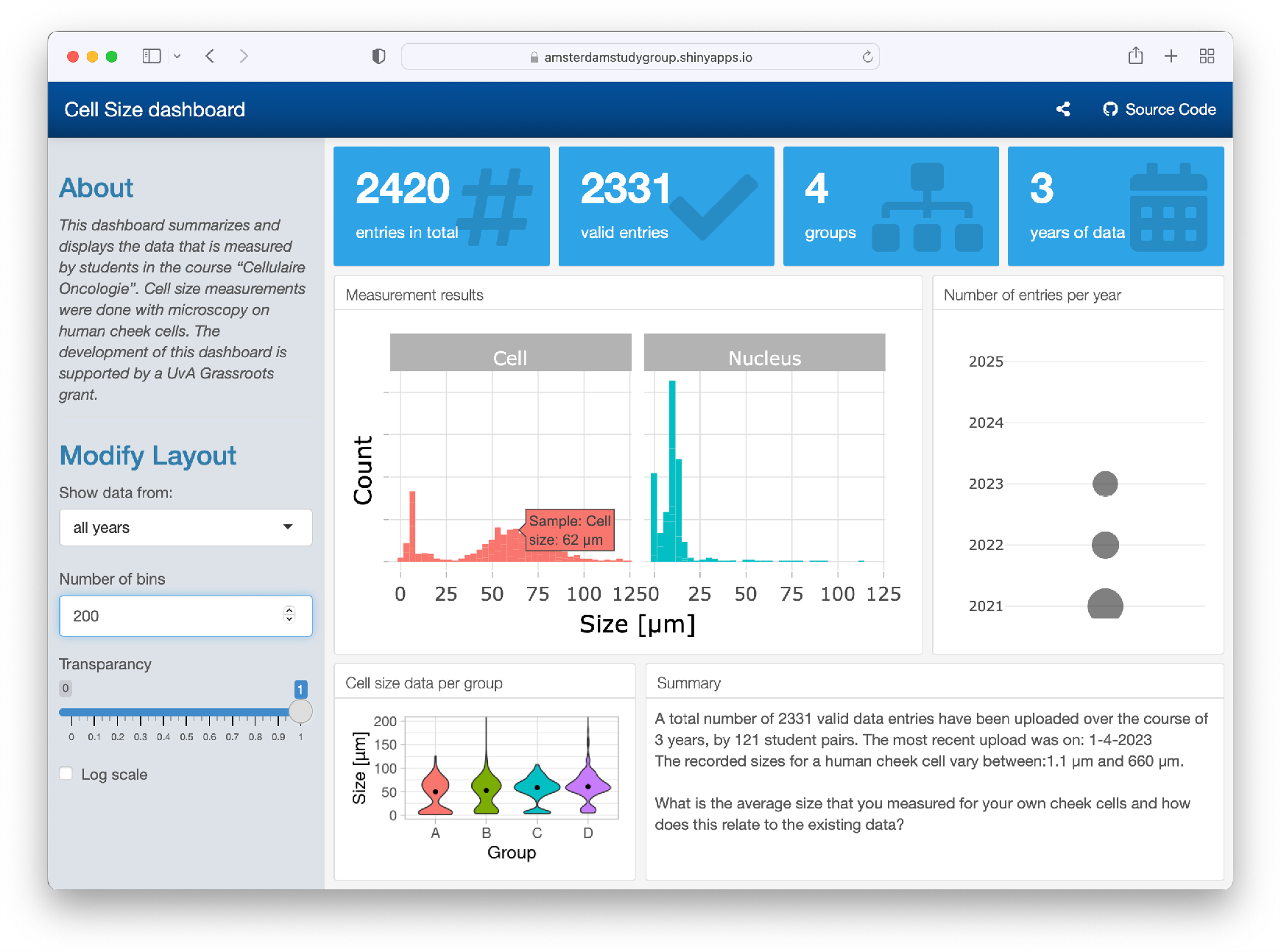
Screenshot of the dashboard that summarized the data on human cheek cell size measurements. The dashboard also displays the total number of entries and other metadata.

The dashboard is interactive and users can select the data from all measurements, or from a single year and the number of bins can be adjusted. Additionally, by hovering over the plots, the values of the data can be read (as shown in Figure 2). The dashboard also shows the data for the 4 different groups and the size distribution of the cells by violin plots.

The histogram on the dashboard is the primary data that is useful for the students. It visualizes the distribution of individual data for both the cell and the nucleus. Since the sizes vary substantially, the data can be shown on a log-scale as well on the dashboard (Figure 3). The main reason that the measurements differ by an order of magnitude, is that the size measurement requires correct calibration of the field of view with a micro ruler. When the calibration is done incorrectly, this will affect the accuracy of the measurement, usually by a factor of 10.

**Figure 3:**
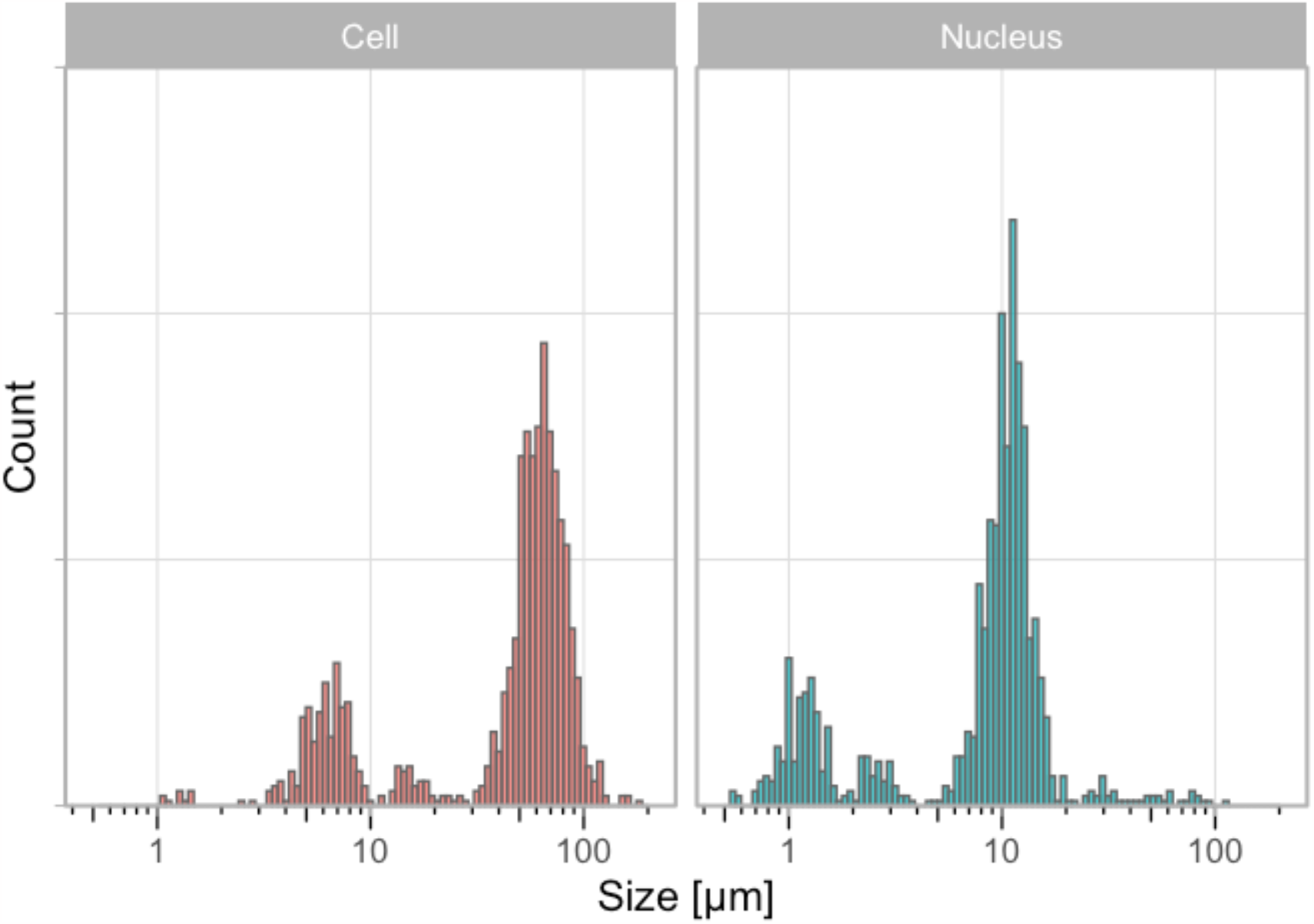
Distribution of the measured size of human cheek cells and their nucleus on a log scale. Aggregated data from three consecutive years (2021-2023). Source: Summarizing the size of cells

Ideally, the students see the multimodal distribution and realize that the peaks are an order of magnitude different. Even if they don’t, they will probably assume that the mode of the distribution is the correct typical value. In any case, it is possible for the students to make a comparison and discuss their results in the context of the historical data.

### Use case 2: Determination of the percentage of cells in S-phase

The aim of the experiment is to determine the percentage of cells that is in the S-phase. To this end, students stain cells that are treated with EdU and DAPI. The images they acquire from both channels are used to quantify the percentage of cells in the S-phase by two analysis methods (manual and semi-automated). The results are uploaded via a Google Form. The collected data can be analysed in multiple ways and here I used it to compare the two analysis methods and, secondly, to obtain an estimate for the percentage of S-phase cells. The data on the two analysis methods, manual and automated, is paired and can be visualised by a doplot in which the pairs of the data are connected (Figure 4). The slopes of the lines vary a lot, whereas the average values per year between the two methods is similar. This implies that there can be substantial differences between the two methods, with roughly a similar number of cases where the automated analysis over-or underestimates the percentage, relative to the manual analysis.

**Figure 4:**
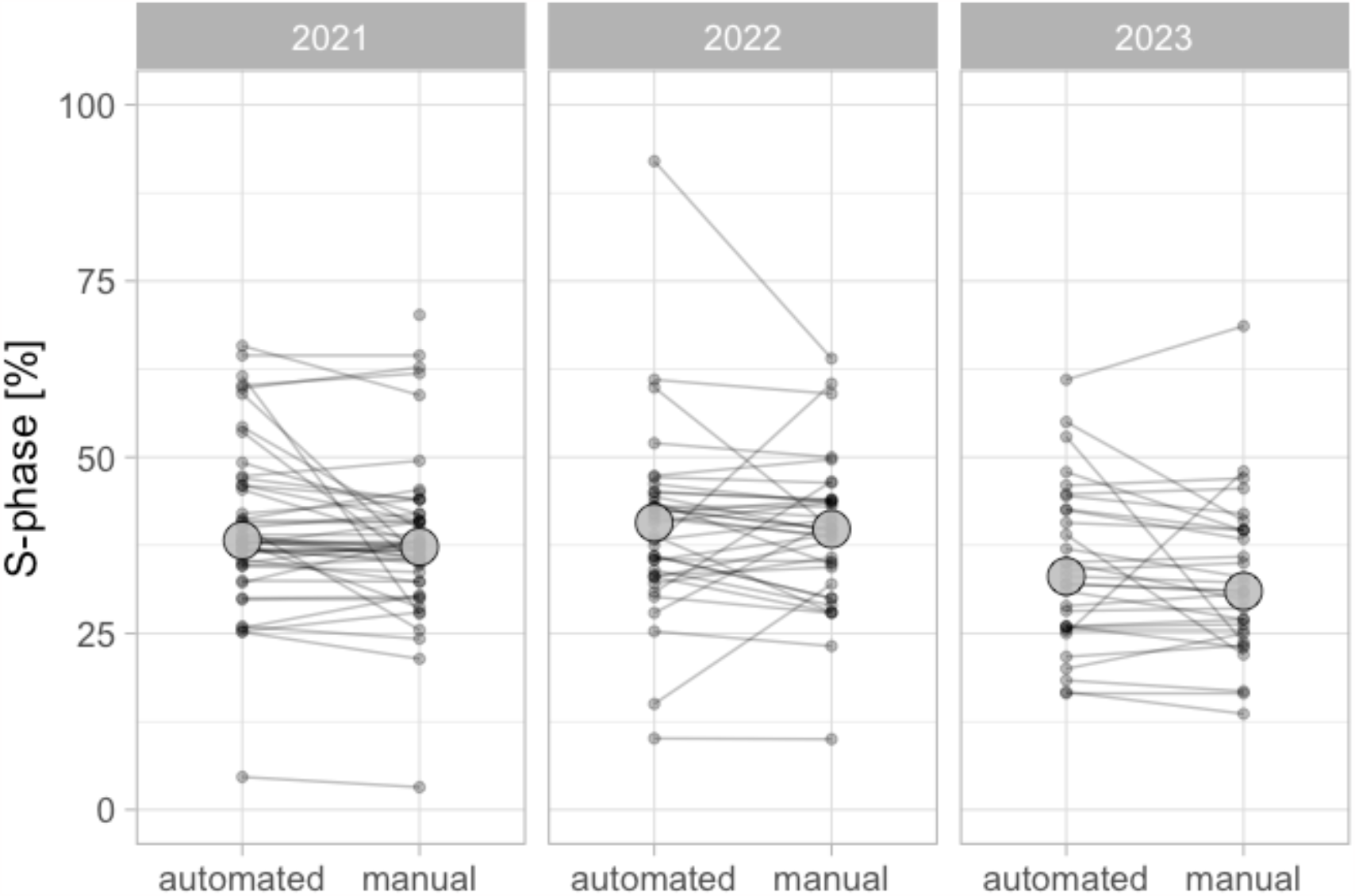
Quantification of the percentage of HeLa cells in the S-phase by EdU incorpotation and fluorescence staining. The data from three different years is shown and a comparison is made between a manual counting method and an automated analysis in ImageJ. The large dot shows the median value, which is comparable between analysis methods. Source: The percentage of cells in the S-phase

There is increasing attention on the role of experimental design in data analysis and visualization. The recently proposed superplot to distinguish biological and technical replicates is an intuitive and straightforward way to communicate the experimental design (Lord et al. 2020). The data on S-phase consists of both technical and biological replicates and is therefore ideally suited to explain the importance of correctly identifying the independent measurements. Here, I treat the data from each group as biological replicate, and the measurements within each group as a technical replicate. The reason is that a group of students stain cells that are from the same passage number and treated at the same time and is therefore a technical replicate. On the other hand, different groups stain different passages of cells and so I treat these as independent observations. When the data is plotted for each individual technical replicate (Figure 5), it can be observed that I received multiple submissions per group, leading to a precise measurement per group. The median values range from 23% to 44%. The average value of the independent observations is 36.7% [N=12, 95%CI: 33.0%-40.3%].

**Figure 5:**
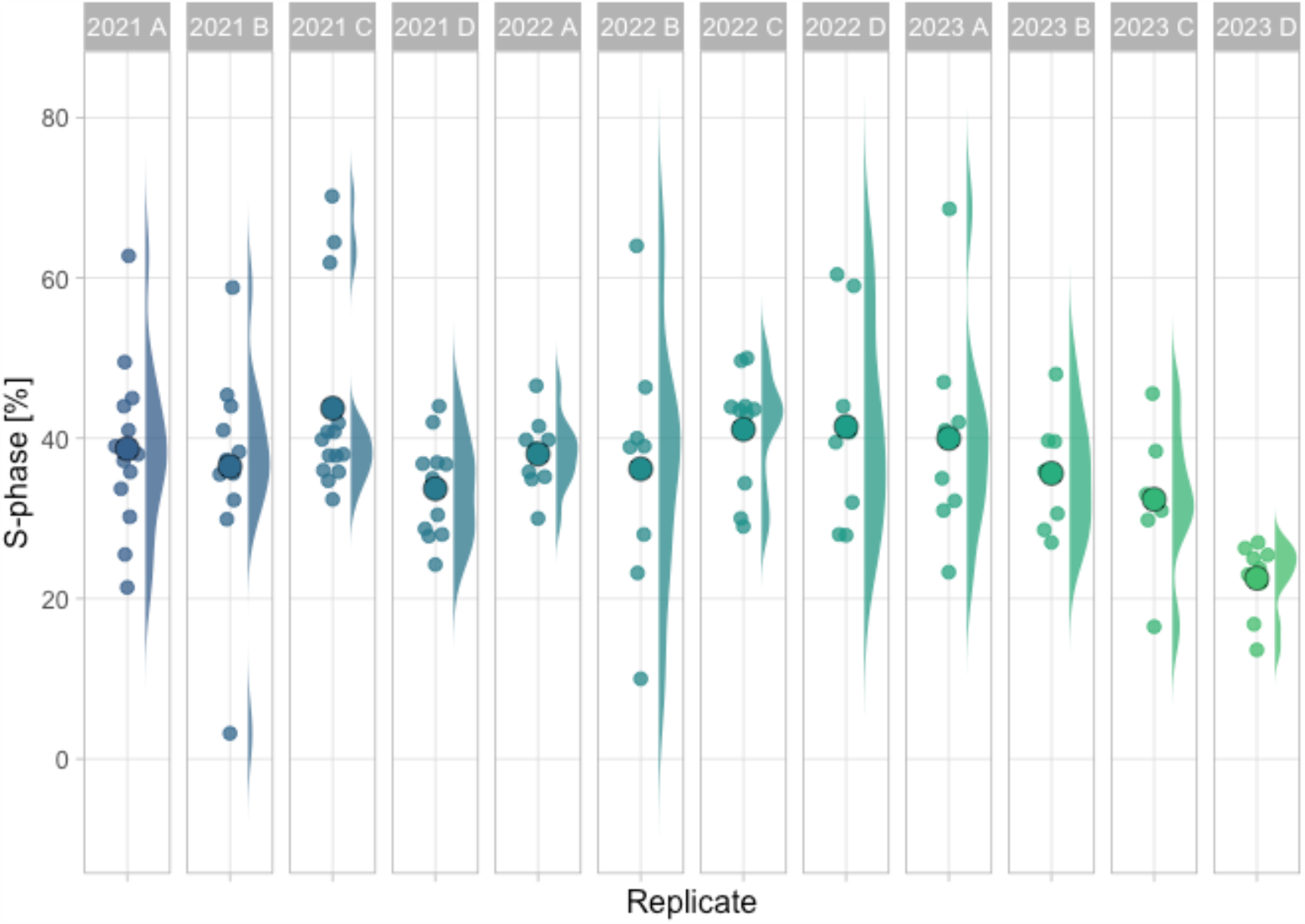
Data on the percentage of cells in e S-phase based on manual analysis. Each group & year defines an independent observations and is shown as dotplot and the distribution. The larger dot reflects the median value. Source: The percentage of cells in the S-phase

## Discussion

Data that is generated in courses is often recorded by individual students or groups of students in reports. However, it can be valuable and interesting to collect, aggregate and (re-)use these data. Here, I present a flexible and straightforward approach to collect and display data from a large group of students and over several years. Here, a combination of Google Forms/Sheets for data collection and R for data processing and visualization is used, and other tools can also be used to achieve data collection and visualization.

In the first use case, the data is displayed in dashboard style. The dashboard gives a quick, interactive and complete overview of the data and it is used by students to evaluate their results. The presentation of the data on the dashboard also acts as a reward for the students as they see that their data has value and is re-used by their peers.

In the second use case, a quarto template is used to process and visualize the data. The use case shows how two analysis methods can be compared and also shows that a high number of truly independent observations can be collected.

For both use cases, the code is available and can be used as a starting point for the processing and visualization of other datasets. This approach is generally applicable and I hope that the use cases provide inspiration for the implementation of studentsourcing in other courses.

In the design of the current course, the groups (A-D) do identical experiments, but it would be straightforward to assign different perturbations (e.g. drug treatments) to different groups and aggregate these data to study effects of perturbations. The perturbations can be done in a blind fashion. After all experiments are completed, a statistical analysis can be performed and the students can discuss the results in their report.

The approach that is presented here is not limited to practical courses. It can also be used to collect data from other crowds, or in collaborative science projects. As such this approach fits in the larger field of citizen science (Silvertown 2009).

Collecting and reusing the data has a number of advantageous aspects. First, a high number of measurements increases the precision of the measurement and therefore allows us to obtain precise numbers. Second, the historical data can be shared with the students and they can interpret and discuss their results in light of the existing data. Third, the obtained data serves as material that can be used to teach data manipulation, statistics and data visualization which is a fundamental aspect of science (Sailem, Cooper, and Bakal 2016). The use cases described in this paper deal with these aspects.

The studentsourcing approach as implemented here has limitations. One limitation is that the outliers or mistakes in the data cannot be traced back to the origin since the data is anonymous. Therefore, providing personal feedback is not possible. Another limitation is that the amount of data that can be uploaded through Google forms is limited. Therefore, uploading of larger datasets (e.g. images), would require a different approach.

An emerging field where a lot of data is required is that of neural networks that are used for artificial intelligence. Particularly the training is resource intensive (Laine et al. 2021) and therefore a studentsourcing approach to distribute the workload would be a potential application.

The aggregation of the data inevitably leads to a discussion on experimental design, as this is important to establish whether measurements are independent or not. This aspect of experimental design has received attention over the last years (Aarts et al. 2015; Sikkel et al. 2017; Eisner 2021) and it is valuable to teach this aspect of data analysis and visualization. Although I have not implemented this yet, I think that having students participate in the data aggregation, creates a very practical opportunity to teach experimental design and the identification of biological units (Lazic, Clarke-Williams, and Munafò 2018). In addition, it may stimulate cooperative learning (Tanner, Chatman, and Allen 2003).

In conclusion, I feel it is valuable to collect data from practical courses and here I report one way to achieve that. I hope that serves as a starting point for others that want to collect, store and use data from large groups of students.

## Data availability

The data is available at: https://doi.org/10.5281/zenodo.8359955

## Code availability

The code for this manuscript is available here: https://github.com/JoachimGoedhart/MS-StudentSourcing and it includes the notebook that was generated to analyze the S-phase data: https://github.com/JoachimGoedhart/MS-StudentSourcing/tree/main/notebooks

The code for the dashboard ‘CellSizeR’ is deposited here: https://github.com/JoachimGoedhart/CellSizeR

## Contributions

J.G. conceived the project, acquired funding, wrote code, and wrote the manuscript.

## Acknowledgments

A blog post by Garrick Aden-Buie was very helpful in the initial phase of this project. Many thanks to the people involved in Quarto, which was used to write and shape this paper. Most importantly, I’d like to thank all students involved in the course that have generously shared their data, making this project a success.

